# Cochlear Innate Immune Homeostasis is altered in the Oncomodulin-Deficient Mouse Model

**DOI:** 10.64898/2026.08.21.745766

**Authors:** Weintari D. Sese, Janith N. Halpage, Mahika V. Palani, Evan J. Paltjon, Kiah C. Sleiman, Aubrey J. Hornak, Dwayne D. Simmons

## Abstract

As part of cochlear innate immunity, cochlear resident macrophages regulate different aspects of tissue maturation, cochlear homeostasis, and injury response. Cochlear resident macrophages exhibit dynamic changes in morphology, distribution, and abundance after cochlear injury. However, in the absence of pathology, regulation of cochlear innate immunity is poorly understood. Since loss of cochlear outer hair cells (OHCs) are indicators of cochlear pathology, we hypothesize that cochlear innate immunity might be sensitive to changes in OHC function. Calcium homeostasis in OHCs is necessary for auditory function, and its dysregulation is associated with hearing loss. However, it is unknown if changes in OHC Ca^2+^ homeostasis are sufficient to alter cochlear innate immunity. Here, we investigate alterations in cochlear innate immunity in a mouse model lacking oncomodulin (OCM), an OHC-specific calcium buffer. Our study focused on the osseous spiral lamina (OSL), a region adjacent to cochlear hair cells. At 1 month, wild-type (WT) mice and *Ocm* knockout (KO) mice have similar hearing thresholds and no evidence of cochlear damage. However, in KO mice, OSL resident macrophages show increased density, altered morphology, and increased spatial segregation closer to the sensory epithelium. Despite these changes in OSL resident macrophages, cytokine profiling revealed no remarkable differences. At 5 months, *Ocm* KO mice show a progressive hearing loss with a frequency dependent loss of OHCs and inner hair cell ribbon synapses, but the density of OSL macrophages remained unchanged. Prior to hearing onset, there was no significant difference in immune cell numbers between *Ocm* WT and KO mice. These findings suggest that cochlear innate immunity is sensitive to OHC calcium buffering following hearing onset.

## Introduction

Acute and chronic insults to the cochlea, such as noise, ototoxic drugs, and aging, induce an innate immune response from cochlear resident macrophages. In addition to an inflammatory response to injury, cochlear resident macrophages participate in cochlear maturation and maintenance of cochlear immune homeostasis under non-pathological conditions (Hirose *et al*., 2005; Okano *et al*., 2008; Hirose *et al*., 2014; Warchol, 2019). Following injury, cochlear resident macrophages participate in the clearance of debris and the recruitment of other effector immune cells to the site of damage through cytokine or chemokine release (Hirose *et al*., 2005; Fujioka *et al*., 2006; Wakabayashi *et al*., 2010; Hirose *et al*., 2017). These macrophages can take on different activation states that involve changes in their abundance, morphology, and expression of immune markers with changes varying between anatomical regions of the cochlea, and mode of injury (Zhang *et al*., 2021). For example, following noise and ototoxic injury, macrophages in the spiral ligament and sensory epithelia take on a pro-inflammatory role that promotes sensory cell degeneration whereas in the spiral ganglion and synaptic area, they have an anti-inflammatory role, promoting neuronal survival and recovery of ribbon synapses (Kaur *et al*., 2015b; Kaur *et al*., 2018; Kaur *et al*., 2019; Manickam *et al*., 2023).

Both acute and chronic insults damage the organ of Corti, the sensory epithelium of the cochlea. The organ of Corti houses two sensory cell types – outer hair cells (OHCs), which amplify and sharpen sound-induced mechanical vibrations through somatic electromotility, and inner hair cells (IHCs), which relay auditory signals to the central nervous system (Ekdale, 2016; Driver & Kelley, 2020). Sensory hair cells initiate transduction of auditory signals to the nervous system through the conversion of sound-induced vibrations to electrical signals in a process called mechanotransduction (MET) (Hudspeth & Jacobs, 1979; Manor & Kachar, 2008; Lelli *et al*., 2009). This process involves the opening of MET channels located on stereocilia and a depolarization response induced by the influx of K^+^ and Ca^2+^ ions (LeMasurier & Gillespie, 2005; Marcotti *et al*., 2014). Although Ca^2+^ makes up a small fraction of the MET current, it serves other important functions such as maintenance of tip link integrity, synaptic transmission, refinement of afferent fiber innervation, and hair cell maintenance and survival (Assad *et al*., 1991; Zhao *et al*., 1996; McPherson, 2018; Ceriani *et al*., 2019).

Noise, ototoxic drugs, and aging target afferent synapses below IHCs and the OHCs. The sensitivity of OHCs to noise and aging is regulated by a small EF-hand Ca^2+^ binding protein, oncomodulin (OCM). In OHCs, OCM is the most abundant calcium binding protein and is localized to compartments associated with calcium dependent processes such as the synaptic zone, along the lateral membrane, and beneath the stereocilia (Sakaguchi *et al*., 1998; Hackney *et al*., 2005; Simmons *et al*., 2010; Tong *et al*., 2016). Studies in *Ocm* knockout (KO) mice show that the absence of this protein leads to early progressive hearing loss and increased susceptibility to noise-induced OHC death (Tong *et al*., 2016; Climer *et al*., 2021; Lachgar-Ruiz *et al*., 2024; Murtha *et al*., 2024; Yang *et al*., 2026). This sensitivity might be tied to Ca^2+^ overload, a common feature of OHC injury, and an impaired ability to buffer intracellular Ca^2+^ (Maurer *et al*., 1993; Fridberger *et al*., 1998; Szucs *et al*., 2006; Zuo *et al*., 2008; Murtha *et al*., 2024). Dysregulated Ca^2+^ homeostasis not only contributes to sensory cell vulnerability and death but may also trigger an inflammatory response within the cochlea (Fridberger *et al*., 1998; Esterberg *et al*., 2013; Esterberg *et al*., 2014; Kurabi *et al*., 2017).

Despite extensive characterization of cochlear macrophages following acoustic or ototoxic injury, how the cochlea responds to sensory cell dysfunction in the absence of damage is not understood. Studies of cochlear innate immunity are mostly focused on models of noise trauma, ototoxicity, or aging, where macrophage recruitment and activation coincide with structural degeneration such as hair cell loss, loss of ribbon synapses or spiral ganglion neuron loss (Fredelius & Rask-Andersen, 1990; Arango Duque & Descoteaux, 2014; Hirose *et al*., 2017; Noble *et al*., 2019; Okayasu *et al*., 2020; Rai *et al*., 2020; Pan *et al*., 2024). However, it is unclear whether changes in cochlear innate immunity occur only in response to tissue damage or can be initiated by earlier sensory cell dysfunction. To address this question, we used a mouse model of OHC dysfunction that gives rise to an accelerated hearing loss phenotype and increased susceptibility to noise damage. Young adult mice retain normal cochlear physiology, thus making this model ideal for distinguishing the impacts of early OHC dysfunction from overt pathology, on the cochlear immune system (Tong *et al*., 2016; Climer *et al*., 2021; Murtha *et al*., 2024).

In this study, we hypothesize that changes in cochlear innate immune sensitivity occur before the onset of structural degeneration, auditory decline, or loss of sensory cells. To test this hypothesis, we characterize cochlear innate immunity in young adult *Ocm* KO mice by assessing macrophage density and activation states before auditory functional decline and after the onset of auditory decline. This study defines how the absence of OCM-mediated calcium buffering in OHCs shapes cochlear innate immunity across young adult, aging and developmental stages. Our findings reveal a previously unrecognized link between calcium dysregulation in OHCs, immune remodeling and auditory decline.

## Methods

### Animals

CBA/CaJ *Ocm^+/+^* (WT) and CBA/CaJ *Ocm^-/-^* (KO) mice of both sexes were used in this study. Samples were obtained before hearing onset at post-natal (P) day 4 – 5, at hearing onset (P12), after hearing onset (1 mo), and at the onset of hearing loss (5 mo) in our *Ocm* KO mouse model. Each experimental group was comprised of a minimum of 6 animals taken from multiple litters. CBA/CaJ *Ocm^-/-^*mice were generated as described in (Climer *et al*., 2021). *Ocm* KO mice were generated from spontaneous germline transmission of the KO allele from the parental line, (C57Bl/6 *Actb^Cre^;Ocm^flox/flox^*) (Tong *et al*., 2016). The original C57Bl/6 *Actb^Cre^;Ocm^flox/flox^*mice were backcrossed onto the CBA/CaJ background to minimize or eliminate the confounding effects of the *Cdh23^ahl^* mutation which is linked to an earlier onset of age-related hearing loss and susceptibility to noise trauma (Davis *et al*., 2001; Noben-Trauth *et al*., 2003). All animals were housed in the Baylor University vivarium and maintained on a 12-hour light/12-hour dark cycle with *ad libitum* access to food and water. All experimental protocols were performed in compliance with the National Institutes of Health (NIH) guidelines and were approved by the Institutional Animal Care and Use Committee at Baylor University.

### Cochlear function assays

Auditory brainstem responses (ABRs) and distortion product otoacoustic emissions (DPOAEs) were used to measure auditory function in adult mice following procedures described in our previous studies (Climer *et al*., 2021; Murtha *et al*., 2024). Mice were anesthetized with an intraperitoneal (i.p) injection of ketamine (100mg/kg) and xylazine (20mg/kg). Body temperature was maintained with a heating pad, and an ophthalmic ointment (OptixCare eye lube) was applied to prevent drying of the eyes as a result of anesthesia. Acoustic stimuli were delivered using a custom-built acoustic assembly in a noise-canceling chamber (Maison *et al*., 2012). ABRs and DPOAEs were recorded from the right ear of each mouse. Both outputs and inputs were processed with a digital I-O board (National Instruments PXI-4461) using the Eaton-Peabody Cochlear Function Test Suite (EPL-CFTS), a LabView-driven data acquisition system.

For ABR measurements, three subcutaneous needle electrodes were placed behind the pinna of the test ear (reference electrode), vertex (active electrode), and above the tail of the mouse (ground electrode). ABR potentials were evoked with a 5 ms tone burst (0.5 ms rise-fall with a cos2 onset, delivered at 35/s). At each sound level, responses were amplified, filtered (100 Hz – 3kHz), and averaged (taken from 512 responses). ABR measurements were obtained at 8, 16, and 32 kHz with sound levels raised in 10-dB steps from 10 to 80 dB. ABR thresholds were defined as the lowest stimulus intensity in which synchronous waveforms with recognizable peaks could be identified.

DPOAEs were measured in response to two primary pure tones, *f1* and *f2* generated by two electrostatic earphones (EC-1, Tucker Davis Technologies, Alachua, FL, USA). Primary tones were presented at 7 frequency pairs, where *f2* = 5.66, 8.0, 11.32, 16.0, 22.56, 32, and 45.2 kHz. Sound levels were increased in 5-dB steps from 10 to 80 dB, and 512 responses were averaged from each sound level. DPOAEs at *2f1-*f2 were recorded in the mouse inner ear canal using a Knowles miniature microphone (Knowles Electronics, Itasca, IL, USA). DPOAE threshold levels for each frequency were determined as the lowest sound pressure level (SPL) at which the distortion product (*2f1-f2*) was consistently above the corresponding noise floor.

### Cochlear harvest and whole mount preparation

Experimental animals’ post-natal day 12 (P12) and above were deeply anesthetized with a lethal dose of sodium pentobarbital (Euthasol®,150mg/kg, i.p.). Following the absence of a toe pinch reflex, mice were transcardially perfused with 1X phosphate buffered saline (PBS) followed by 4% (w/v) paraformaldehyde (PFA) [EMS, EM Grade, in 1X PBS]. Inner ears were dissected from the temporal bone and an insect pin (FST, size 000), was used to open a small hole in the apex. The cochlea was perfused with 4% PFA through the round and oval windows. P4 – P5 pups were euthanized by placing on ice until unresponsive. Temporal bones were harvested and cochlear spirals carefully obtained.

For fluorescent immunolabelling with synaptic markers, left inner ears were post-fixed for 20 minutes while right inner ears, used for all other immunolabelling procedures, were post-fixed overnight. Following fixation, inner ears were washed with 1X PBS and decalcified at room temperature for 2 – 4 days. For whole mount dissections, organs of Corti were isolated by carefully trimming away the spiral ligament and modiolar nerve stump. Microdissected pieces spanning all cochlear frequency regions from base to apex were obtained.

### mRNA expression analysis

Cochleae from both ears were quickly harvested in ice-cold 1X Dulbecco’s phosphate-buffered saline (DPBS; Gibco, 14190094). After harvest, cochleae were transferred to a petri dish containing RNAlater™ solution (Invitrogen, AM7020) and the bony shell chipped away. Subsequently, cochlear tissue were transferred to a lysis buffer, Buffer RLT Plus, and homogenized using a handheld homogenizer. Total RNA was extracted following manufacturer’s instructions using the RNeasy plus Micro kit (Qiagen) and was reverse transcribed using the iScript advanced cDNA reverse transcription kit (Bio-Rad).

qRT-PCR was performed on a CFX96 Real time system (Bio-Rad) using the SYBR Green PCR Master Mix Kit (Bio-Rad). The cycle threshold (Ct) values of the target genes were first normalized to the average expression level of *Gapdh* to generate the ΔCt values. Analysis of relative gene expression levels between *Ocm* WT and KO cochleae was completed using the 2^-ΔΔCt^ method as previously reported (Livak & Schmittgen, 2001) with ΔΔCt values normalized to the average expression level of the target gene in *Ocm* WT cochlea.

Primers used are as follows: *Gapdh* forward 5’-AGA CAG CCG CAT CTT CTT GT-3’ and reverse 5’-CTT GCC GTG GGT AGA GTC AT-3’; *Il-6* forward 5’-CTT CAC AAG TCG GAG GCT TAA T-3’ and reverse 5’-ACACTGGGTCTTCATCAGTTTC-3’; *Il-1*β forward 5’-ATG GGC AAC CAC TTA CCT ATT T-3’ and reverse 5’-GTT CTA GAG AGT GCT GCC TAA TG-3’; *Il-4* forward 5’-TTT GCG TGG TTT CTA GGG ATA G-3’ and reverse 5’-GCT GAG GCA TGG ATC TGT TAG-3’; *Il-10* forward 5’-CTC TTC CTC CTC CTT CTC TTC T-3’ and reverse 5’-GGG TAA TAG GTG CTG GAA ATA GG -3’.

### Immunocytochemistry

Immunofluorescence staining was performed using procedures described in our previous publications(Yang *et al*., 2023; Murtha *et al*., 2024). Briefly, microdissected pieces were incubated in 30% fresh sucrose (in PBS) for 15 minutes on a shaker at room temperature, frozen at -80 until frozen through, then thawed at 37. Pieces were rinsed in PBS 3 times and incubated at room temperature for 1 hr in permeabilization and blocking solution [5% normal horse serum (Sigma-Aldrich, H0146), 0.3% Triton X-100 (Bio-Rad, 1610407) in PBS (5% NHS-T)]. Microdissected pieces were incubated overnight at 37 in primary antibodies diluted in 1% NHST. For synaptic immunolabelling, the following primary antibodies were used: mouse CtBP2 (BD Biosciences, 612044, 1:200), mouse GluR2 (Millipore, MAB397, 1:200), rabbit Myo7a (Proteus Bio, 25-6790, 1:200). For immune cell immunolabelling, microdissected pieces were incubated with the following primary antibodies: goat CD45 (R & D systems, AF114, 1:200), rabbit IBA1 (Wako Fujifilm, 019-9714, 1:200) and rat CD68 (Bio-Rad, MCA1957GA, 1:50). Following overnight incubation in primary antibodies, pieces were rinsed 6 times in PBS and incubated in the dark with the appropriate species-specific Alexa Fluor secondary antibodies and phalloidin (diluted in 1% NHST) for 2 hrs at 37. Pieces were rinsed 3 times in PBS and incubated in Hoechst 33342™ (Invitrogen™, H3570, 1:7500) for 5 minutes at room temperature prior to the final rinse. Microdissected pieces were mounted with VECTASHIELD® mounting media (Vector laboratories©) onto glass slides, coverslipped and sealed with clear nail polish. Cohorts of samples were immunostained at the same time to allow for direct comparisons.

### Confocal Microscopy

All images were acquired on a Zeiss LSM800 confocal laser scanning microscope. Low-power images of cochlear microdissected pieces were obtained with a 10x air objective (0.3 NA) and cochlear frequency maps generated using ImageJ (version 2.16.0, Fiji, NIH, Bethesda) using the *Measure_line.class* plug-in (downloaded from the Massachusetts Eye and Ear Histology Core Website). Subsequently, high resolution confocal z-stack images corresponding to the apical (5.6, 8 and 11.3 kHz), middle (16 kHz and 22.6 kHz) and basal frequency regions (32 kHz and 45.2 kHz) were acquired using the generated frequency map.

### Mouse cytokine array analysis

The proteome profiler mouse cytokine array kit (R & D systems, ARY006) was used for the simultaneous detection of 40 mouse cytokines. Whole cochlear lysates were prepared according to the manufacturer’s instruction. A sample was generated by pooling four cochleae and protein quantified using the Pierce™ Bicinchoninic Assay (BCA) protein assay kit (ThermoFisher Scientific, 23225). A total protein amount of 250 µg was used for the array. The chemiluminescent signal on each membrane was collected using a ChemiDoc imaging system (Bio-Rad). Mean grey values were collected for each spot on the array using ImageJ and corrected for background intensity. Spots were then normalized to the mean grey values of positive reference spots on the same membrane. Finally, fold changes were quantified by normalizing the mean grey values of *Ocm* KO spots to the corresponding spot on the WT membrane.

### Synapse and OHC counts

For IHC synaptic counts, high resolution confocal z-stacks of hair cells and synapses were acquired using an oil-immersion objective (Plan-Apochromat 63x, 1.4 NA). Acquired z-stack images spanned the entire synaptic pole of 12 – 14 IHCs (identified by Myo7a immunolabeling). Analysis of synapses (juxtaposed CtBP2 and GluR2 puncta) and orphan ribbons (CtBP2 puncta) were performed in Imaris (Oxford Instruments). Threshold adjustments for all channels were done to reduce any background while not compromising actual signal. A surface of the IHC region was created using the surfaces tool of Imaris, and the CtBP2 and GluR2 channels were subsequently masked. To determine the number of ribbon synapses and orphan ribbons, the spot detection feature of Imaris was used. Briefly the size of CtBP2 and GluR2 puncta was estimated by measuring the XY diameter of multiple puncta and getting an average value. This value was then used to isolate individual CtBP2 and GluR2 puncta. Finally, a filter was applied for distance to nearest spots for CtBP2 and GluR2 to determine colocalization. The minimum possible colocalization was defined as the summed XY diameter of CtBP2 and GluR2 puncta. CtBP2, GluR2 and juxtaposed CtBP2-GluR2 spots were manually confirmed. Number of ribbon synapses and orphan ribbons per IHC was determined by dividing the total number of CtBP2-GluR2 and CtBP2 puncta by the total number of IHCs present.

For OHC counts, images encompassing all frequency regions were acquired using an air objective lens (Plan-Apochromat 20x, 0.6 NA). OHCs were manually counted using the ImageJ cell counter plug-in and represented as OHC counts per image field.

### Macrophage density and morphological analysis

Macrophages were identified by positive CD45 and Iba1 immunolabeling, and high-resolution z-stacks were acquired with a 10x air objective (0.3 NA). Z-stack images were acquired such that start and stop points were defined as when macrophages came into focus and out of focus, respectively. This was done to ensure the entire depth of cells were captured for subsequent morphological analysis. Macrophage density was determined in the osseous spiral lamina (OSL) only by manual counting using the ImageJ cell counter plug-in. Results were reported as the number of macrophages normalized to an OSL area of 0.1mm^2^. Actual OSL areas were determined by manually creating a surface using the “surfaces” feature of Imaris. The “surfaces” feature of Imaris was also used for the 3D reconstruction of individual macrophages. Representative OSL images shown here were acquired with a 20x objective lens (0.6 NA).

“MotiQ”, an open-source toolbox available as ImageJ plug-ins was used for all morphological analysis of macrophages (Hansen *et al*., 2022). The workflow for analysis involved a three-step process using the following plug-ins: *MotiQ cropper*, *MotiQ thresholder* and *MotiQ 2D analyser*. Briefly, MotiQ cropper was used to select individual macrophages in our region of interest. The generated images of individual macrophages were subjected to image segmentation using the *MotiQ thresholder*. Finally, *MotiQ 2D analyser* was used to quantify morphological parameters such as cell size, tree length, ramification index and spanned area. A minimum of 10 randomly selected macrophages were analyzed from 2 cochleae per group.

Spatial distribution of macrophages was determined using the “analyze particles” plug-in of ImageJ. X and Y centroid values were determined for each macrophage in 20x images of the OSL. Briefly, images were converted to 8 bit and contrast was adjusted as necessary. The images were made binary and threshold adjusted as necessary. Following threshold adjustment, the “fill holes” feature was used to fill empty areas detected within each macrophage. Finally, X and Y centroid values were determined using the “analyze particles” plugin by setting a pixel size as appropriate to exclude any particles other than macrophages.

### Statistics

Statistical analyses were carried out using GraphPad Prism 10 (GraphPad, San Diego, CA). An unpaired *t*-test was used to compare means between two groups. Comparisons of multiple groups were conducted using one-way or two-way ANOVA followed by Bonferroni’s post-hoc test unless otherwise stated in the figure legends. Data is presented as means ± standard deviation (SD). Statistical significance is denoted with asterisks as the following: * *p < 0.05,* ** *p < 0.01,* *** *p < 0.001,* **** *p < 0.0001*. Lack of statistical significance is denoted as “ns” where *p > 0.05*.

## Results

### *Ocm* KO mice have increased macrophage density before the onset of hearing loss

Increased macrophage numbers and alterations in morphology are associated with cochlear injury (Hirose *et al*., 2005; Sato *et al*., 2010; Frye *et al*., 2017). The recruitment and activation of macrophages following cochlear injury is associated with hair cell death and synaptic loss, with several studies implicating macrophages in debris clearance and repair of ribbon synapses (Kaur *et al*., 2015a; Kaur *et al*., 2015b; Kaur *et al*., 2018; Kaur *et al*., 2019; Manickam *et al*., 2023). In this study, we evaluate cochlear macrophage density and morphology in the *Ocm* KO mouse, which have a normal hearing phenotype in the young adult but an early progressive hearing loss and increased susceptibility to noise trauma (Climer *et al*., 2021; Murtha *et al*., 2024; Yang *et al*., 2026).

Our analyses were restricted to the osseous spiral lamina (OSL), which contains more than half the population of cochlear resident macrophages, thus making it a representative region for assessing cochlear macrophage density and morphology (Okano *et al*., 2008). We hypothesize that the absence of OCM in OHCs might lead to an altered immune state in young adults, which may contribute to increased susceptibility to hearing loss. We immunostained cochlear whole mounts from 1-month old *Ocm* KO (*Ocm^-/-^*) and age-matched *Ocm* WT (*Ocm^+/+^*) mice with CD45, a pan-leukocyte marker, and IBA1, a macrophage and microglia-specific marker **(Figure 1A & B)**. Quantitative analyses revealed significantly higher macrophage densities in the OSL of *Ocm* KO cochleae across all frequency regions (apex: *Ocm* WT = 24.207 ± 3.224, *Ocm* KO = 27.865 ± 2.125, middle: *Ocm* WT = 26.225 ± 2.114, *Ocm* KO = 28.650 ± 2.778, base: *Ocm* WT = 26.733 ± 1.841, *Ocm* KO = 32.142 ± 3.971, *p* < 0.001, two-way ANOVA) **(Figure 1E)**.

**Figure 1.**
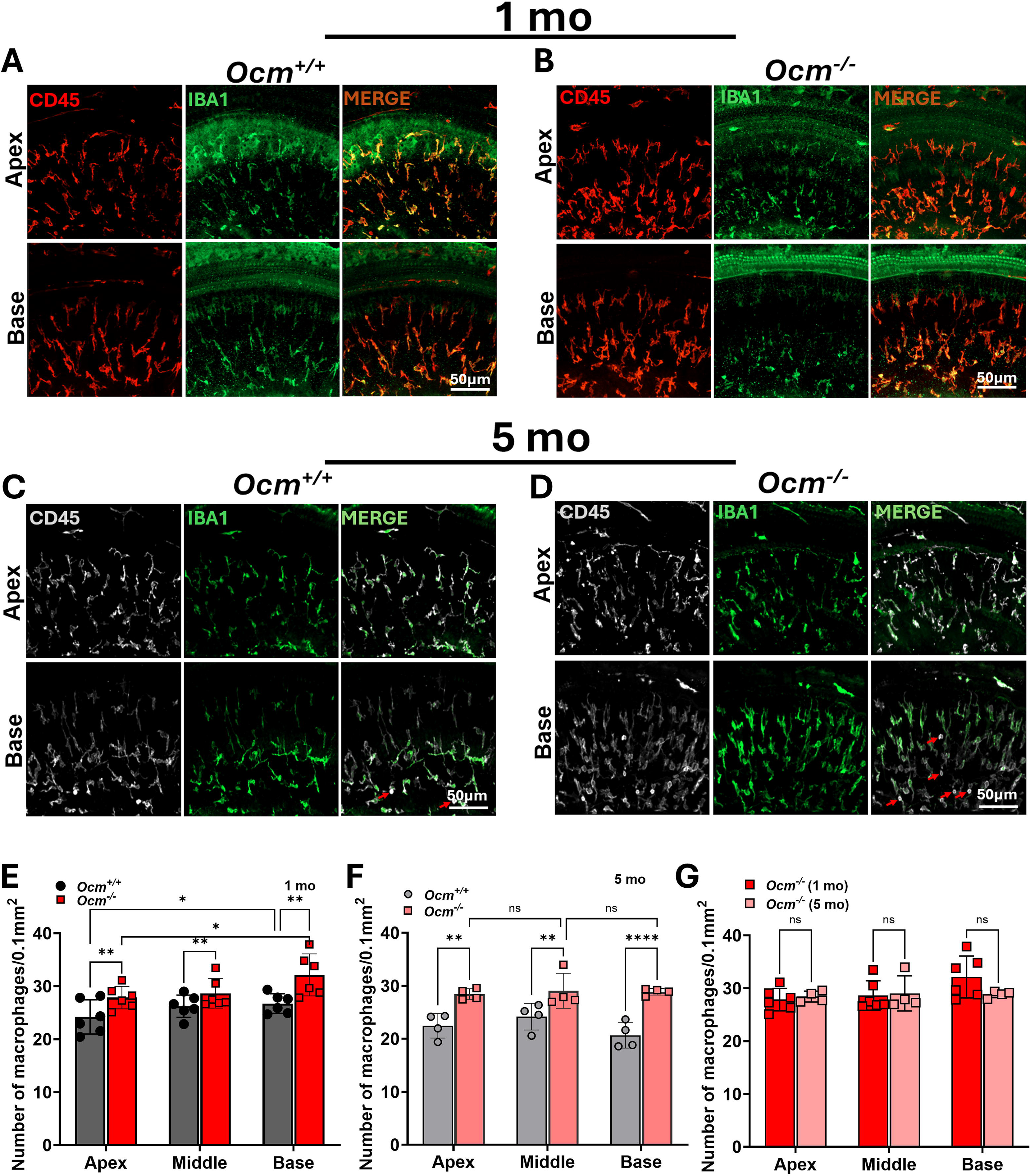
*Ocm* KO cochleae have increased macrophage density compared to WT cochleae. (A, **B**) Representative images of immune cells present in the osseous spiral lamina (OSL) of apical (5 – 11 kHz) and basal (32 – 45 kHz) frequency regions in young adult *Ocm* WT **(A)** and KO **(B)** cochleae. Immune cells are labeled with the pan-leukocyte marker, CD45 (red) and the macrophage-specific marker, IBA1 (green). Scale bar = 50 µm. **(C, D)** Representative images of CD45 (white) and IBA1 (green) immunolabeling in the OSL of 5 month old *Ocm* WT **(C)** and *Ocm* KO **(D)** cochleae. **(E & F)** Quantification of OSL macrophages across frequency regions in young adult **(E)** and aged **(F)** *Ocm* WT and KO cochleae. **(G)** Quantification of OSL macrophages across frequency regions in young adult and aged *Ocm* KO cochleae only (n = 4 to 6 cochleae per experimental group). Data are presented as mean ± SD, two-way ANOVA with Bonferroni’s post hoc. * *p < 0.05;* ** *p < 0.01;* **** *p < 0.0001*; ns, not significant

To determine if the changes in macrophage density seen in young adult *Ocm* KO mice persisted after the onset of hearing loss in these mice, we immunostained for macrophages in 5 mo old WT and KO mice **(Figure 1C & D)**. While there was an increase in macrophage density compared to WT cochleae (apex: *Ocm* WT = 22.455 ± 2.308, *Ocm* KO = 28.438 ± 1.025, *p* < 0.01, middle: *Ocm* WT = 24.173 ± 2.514, *Ocm* KO = 29.040 ± 3.315, *p* < 0.01, base: *Ocm* WT = 20.680 ± 2.424, *Ocm* KO = 28.858 ± 0.626, *p* < 0.0001; two-way ANOVA) **(Figure 1F)**, there was no difference between macrophage density in young adult *Ocm* KO mice when compared to aged KO mice **(Figure 1G)**. Additionally, we observed the presence of a small number of CD45 positive but IBA1 negative cells (indicated by red arrows in **Figure 1C & D**) suggestive of non-resident immune cell populations in both aged *Ocm* WT and KO OSL.

Together, these results suggest that OHC dysfunction resulting from impaired Ca^2+^ homeostasis is sufficient to alter the cochlear innate immunity in the absence of damage to cochlear structures. While reports suggest that macrophage density is highest in regions undergoing active sensory cell degeneration (Frye *et al*., 2017), the lack of correlation between OHC loss and OSL macrophage density in aged *Ocm* KO mice suggests cues other than sensory cell degeneration in the recruitment of macrophages to damage sites.

### Macrophage morphology, spatial distribution and activation state in the OSL varies between young adult *Ocm* WT and KO cochlea

Previous studies indicate that macrophage morphology changes with cochlear location and damage (Yang *et al*., 2015; Okayasu *et al*., 2020; Pan *et al*., 2024) **(Figure 2A)**. To determine if the increased macrophage density in the OSL of young adult *Ocm* KO cochlea accompanied alterations in macrophage morphologies similar to changes described, we assessed the morphologies of OSL resident macrophages. We quantified metrics such as cell size, spanned area and ramification index which are commonly used to describe the morphology and activation states of microglia, the resident macrophages of the central nervous system (Fernández-Arjona *et al*., 2017; Hansen *et al*., 2022).

**Figure 2.**
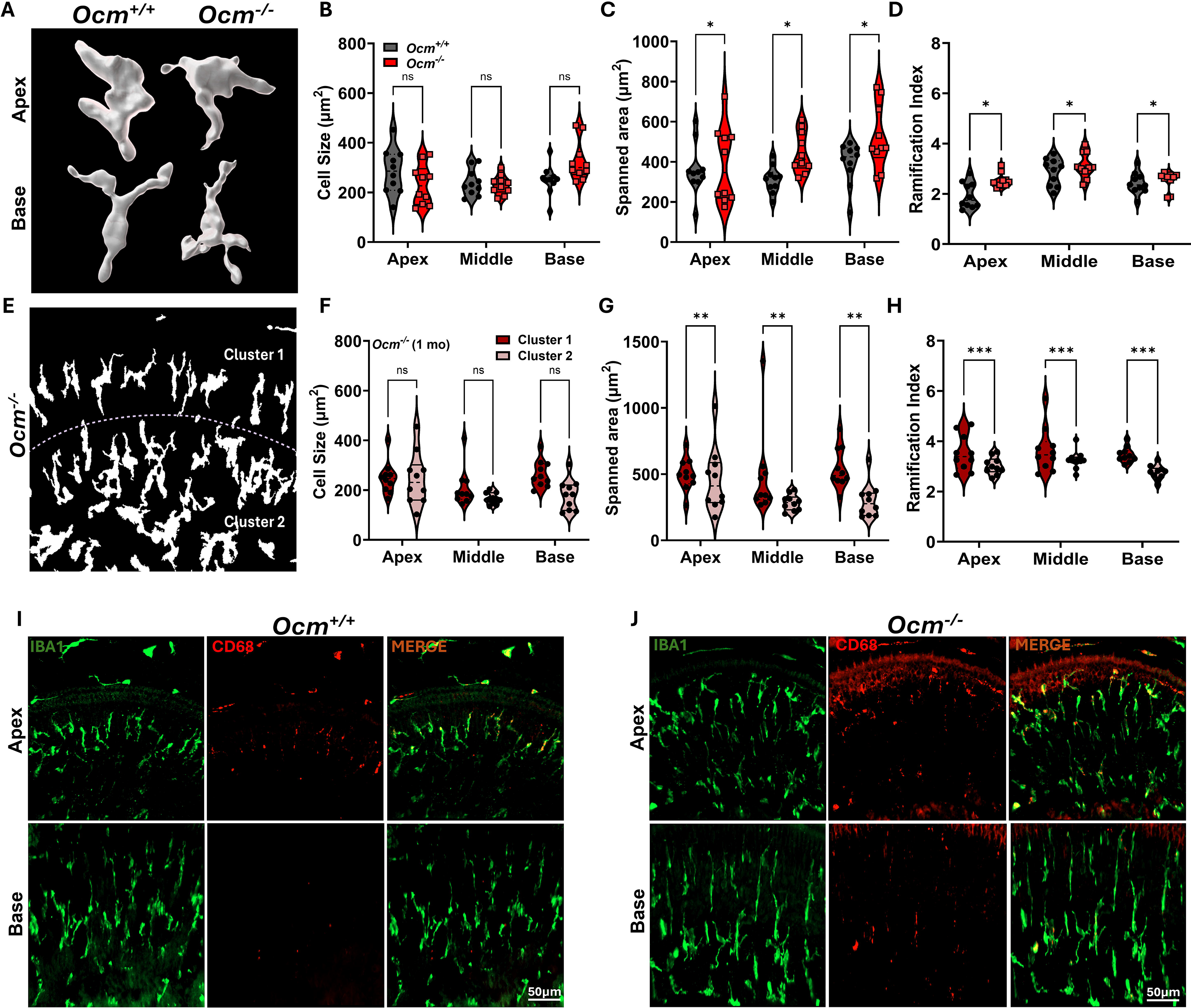
Macrophage morphology, spatial distribution and activation states are altered in the OSL of *Ocm* KO cochleae. **(A)** 3D reconstructions of representative OSL macrophages in the apical and basal regions of *Ocm* WT (left panel) and *Ocm* KO (right panel) cochleae. **(B – D)** Quantitative analysis of macrophage morphology: cell size **(B)**, spanned area **(C)** and ramification index **(D)** in young adult and *Ocm* WT and KO cochleae. **(E)** Binarized image showing a distinct spatial distribution of OSL macrophages in *Ocm* KO cochleae. **(F – H)** Quantitative analysis of cluster 1 and cluster 2 OSL macrophages in young adult *Ocm* KO cochlea: cell size **(F),** spanned area **(G),** and ramification index **(H)** (n ≥ 10 macrophages per group). **(I & J)** Representative images of macrophages labeled with the macrophage specific marker, IBA1 (green) and a marker of activated macrophages, CD68 (red) in young adult *Ocm* WT **(I)** and KO **(J)** cochlea. Data are presented as mean ± SD, two-way ANOVA with Bonferroni’s post hoc. * *p < 0.05;* ** *p < 0.01;* *** *p < 0.001*; ns, not significant

While macrophage cell size across frequency regions was not significantly different between *Ocm* WT and *Ocm* KO cochlea (*p* > 0.05, two-way ANOVA), parameters such as spanned area and ramification index were significantly higher (*p* < 0.05, two-way ANOVA) in OSL macrophages of *Ocm* KO cochleae. (**Figure 2B – D)**. These morphological features are characteristic of macrophages in a heightened surveillance state, suggesting that OSL macrophages in *Ocm* KO cochlea exhibited enhanced surveillance of their microenvironment even under homeostatic conditions.

Furthermore, in some *Ocm* KO cochleae, we observed a distinct distribution of macrophages in the OSL, which was confirmed by an unbiased, spatial distribution analysis based on X and Y centroid coordinates of individual macrophages **(Figure S1A – F)**. Based on spatial distribution, macrophages were classified into two clusters - cluster 1 macrophages with lower Y centroid values and located closer to the IHC region, and cluster 2 macrophages located closer to the spiral ganglion neurons **(Figure S1A, S1D, 2E)**. Morphometric analyses revealed that macrophages in cluster 1 and cluster 2 had comparable cell sizes (*p* > 0.05, two-way ANOVA) **(Figure 2F)**. Morphological parameters such as spanned area, and ramification index were significantly higher in cluster 1 macrophages compared to cluster 2 macrophages across all frequencies (spanned area, *p* < 0.01, ramification index, *p* < 0.0001, two-way ANOVA) **(Figure 2G & H)**. Again, these features are consistent with increased surveillance activity suggesting that OSL macrophages closer to the IHC region demonstrate heightened surveillance.

Considering the changes in resident macrophage density and morphology observed in young adult *Ocm* KO cochleae, we sought to determine if these changes accompanied changes in macrophage activation states. We analyzed the expression of CD68, a lysosomal marker associated with increased macrophage phagocytosis and recruitment (Holness *et al*., 1993; Gawande *et al*., 2025). *Ocm* KO cochleae seemed to have a greater number of CD68 expressing macrophages in basal frequency regions compared to WT cochlea while CD68 expression was comparable in apical frequency regions between both genotypes **(Figure 2I & J)**. The increased expression of CD68 in the base may indicate changes in macrophage states that can lead to vulnerability to hearing loss in *Ocm* KO mouse model.

### Altered Cochlear Innate Immunity is independent of changes in ABRs and degeneration of ribbon synapses

Our previous work demonstrates that OCM is necessary for maintaining auditory function and its absence is correlated with early, progressive hearing loss, and an increased vulnerability to noise (Tong *et al*., 2016; Climer *et al*., 2021; Murtha *et al*., 2024; Yang *et al*., 2026). In the current study, we evaluated if the observed changes in macrophage density and morphology in 3 – 4 wks old *Ocm* KO occur independently of any changes in auditory function. We evaluated ABRs at 8, 16 and 32 kHz in *Ocm* WT and *Ocm* KO mice. *Ocm* KO mice had ABR threshold levels comparable to WT mice (*p* > 0.05, two-way ANOVA) **(Figure 3A)**. To assess auditory nerve function, ABR wave I amplitude and latency was evaluated at 8 kHz and 32 kHz. At suprathreshold levels wave 1 amplitude and latency were not significantly different in *Ocm* WT and *Ocm* KO mice at 8 kHz (*p* > 0.05, two-tailed, unpaired *t*-test) **(Figure 3B)**, and 32 kHz (*p* > 0.05, two-tailed, unpaired *t*-test) **(Figure 3C)**.

**Figure 3.**
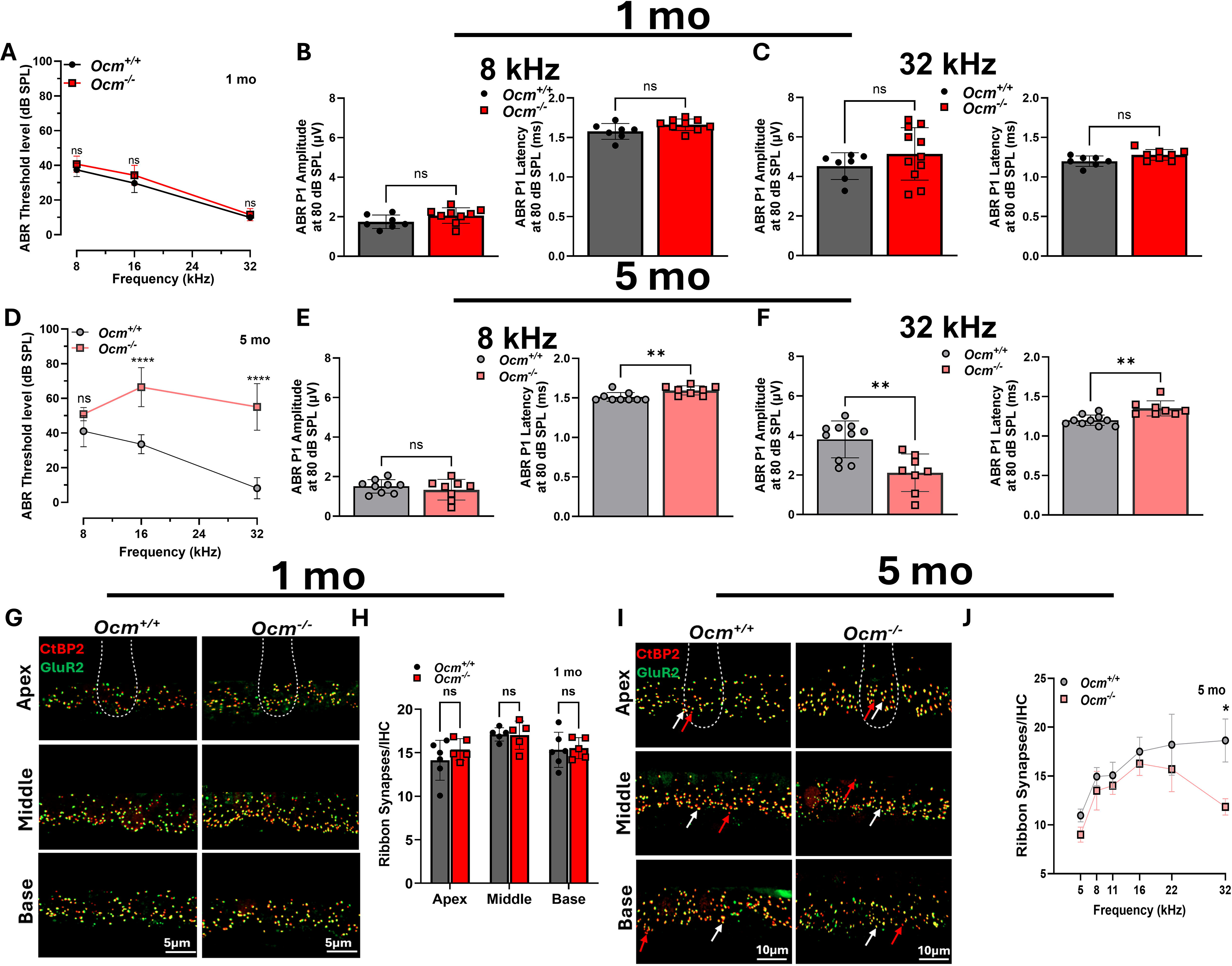
Auditory Brainstem Responses (ABRs) and ribbon synaptic density change with age in *Ocm* KO mice. (A –. **C)** ABR threshold levels **(A)** and wave 1 amplitude (left) and latency (right) at 8 kHz **(B)** and 32 kHz **(C)** in young adult *Ocm* WT and KO mice. **(D – F)** ABR threshold levels **(D)** ABR wave 1 amplitude (left) and latency (right) at 8 kHz **(E)** and 32 kHz **(F)** in aged *Ocm* WT and KO mice. **(G & I)** Representative images of IHC ribbon synapses in apical (5 – 11 kHz), middle (16 – 22 kHz) and basal (32 – 45 kHz) frequency regions in young adult *Ocm* WT and KO cochleae **(G)** and aged *Ocm* WT and KO cochleae **(I)**. Presynaptic ribbons were labeled with CtBP2 (red), and post-synaptic densities with GluR2 (green). Scale bar = 5 µm (young adult) and 10 µm (aged). Red arrows point to orphan ribbons (CtBP2 only) and white arrows point to ribbons synapses (juxtaposed CtBP2 and GluR2 puncta) **(H & J)** Quantification of ribbon synapses across frequency regions in young adult **(H)** and aged **(J)** *Ocm* WT and KO cochleae. Data are presented as mean ± SD, n = 6 to 11 mice per experimental group. Statistical analysis was determined using two-way ANOVA with Bonferroni’s post hoc **(D, H & J)** and unpaired *t*-test **(B, C, E & F)**. * *p < 0.05;* ** *p < 0.01;* **** *p < 0.0001*; ns, not significant

In contrast, aged *Ocm* KO mice exhibited significantly elevated ABR threshold levels at 16 kHz (*Ocm* WT = 33.527 ± 5.476 dB SPL, *Ocm* KO = 66.453 ± 11.253 dB SPL, *p <* 0.0001, two-way ANOVA) and 32 kHz (*Ocm* WT = 8.185 ± 6.072 dB SPL, *Ocm* KO = 55.081 ± 13.459 dB SPL, *p <* 0.0001, two-way ANOVA) **(Figure 3D).** *Ocm* KO mice had unchanged amplitudes at 8 kHz but significantly lower amplitudes at 32 kHz (*Ocm* WT = 3.804 ± 0.934 µv, *Ocm* KO = 2.114 ± 0.949 µv, *p <* 0.01; two-tailed, unpaired *t*-test) with delayed latencies (8 kHz, *Ocm* WT = 1.511 ± 0.0056 ms, *Ocm* KO = 1.595 ± 0.058 ms, *p <* 0.01, 32 kHz, *Ocm* WT = 1.200 ± 0.065 ms, *Ocm* KO = 1.350 ± 0.095 ms, *p <* 0.01; two-tailed, unpaired *t*-test) **(Figure 3E – F).**

Changes in macrophage density and morphology are often associated with cochlear injury, which causes loss of OHCs and ribbon synapses (Kaur *et al*., 2015b; Frye *et al*., 2017; Frye *et al*., 2019). These changes have been shown to be beneficial or detrimental depending on the mode of injury (Manickam *et al*., 2023; Pan *et al*., 2024; Sung *et al*., 2024). In any case, changes in macrophage density and morphology can be either a cause or effect of hair cell and/or synaptic loss. While we observed changes in macrophage density and morphology in young adult *Ocm* KO, these changes were independent of ribbon synapse loss (*p* > 0.05, two-way ANOVA) **(Figure 3G & H)**. In aged *Ocm* KO cochleae, there was a trend of reduced ribbon synapses across all frequency regions, but this difference was significant only at 32 kHz (*Ocm* WT = 18.640 ± 2.206, *Ocm* KO = 11.845 ± 0.836, *p* < 0.05, two-way ANOVA) **(Figure 3I & J).**

These results indicate that young adult *Ocm* KO mice do not exhibit any auditory nerve dysfunction despite an increase in macrophage density and differences in morphology. While macrophage numbers increased relative to age matched WT mice but unchanged relative to aged *Ocm* KO mice, these mice displayed elevated ABR threshold levels and loss of ribbon synapses. These findings suggest that the disruption of OHC calcium homeostasis causes an altered immune state in the OSL that precedes synaptic degeneration and elevation of ABR thresholds.

### Changes in Cochlear Innate Immunity occur before OHC dysfunction in *Ocm* KO mice

Changes in cochlear innate immunity are associated with impaired functioning and loss of OHCs after injury. Taking into consideration the observed changes in macrophage density and morphology in young adult mice, we sought to determine if there were any changes in OHC function. To assess OHC function, DPOAE responses were evaluated. There was no significant difference in DPOAE threshold levels in young adult *Ocm* KO mice when compared to WT mice (*p* > 0.05, two-way ANOVA) **(Figure 4A).** Similar to DPOAE threshold levels, no statistically significant difference was observed in DPOAE I/O responses at 8 kHz and 32 kHz **(Figure 4B & C)**. Similar to our previous studies, older *Ocm* KO mice demonstrated elevated DPOAE threshold levels at the mid to high frequencies (16 kHz, *Ocm* WT = 28.531 ± 4.246 dB SPL, *Ocm* KO = 56.800 ± 14.589 dB SPL, *p <* 0.0001, 22 kHz, *Ocm* WT = 32.825 ± 8.342 dB SPL, *Ocm* KO = 79.049 ± 1.534 dB SPL, *p <* 0.0001, 32 kHz, *Ocm* WT = 26.197 ± 5.810 dB SPL, *Ocm* KO = 78.399 ± 3.390 dB SPL, *p <* 0.0001, and 45 kHz, *Ocm* WT = 36.192 ± 9.186 dB SPL, *Ocm* KO = 76.721 ± 2.156 dB SPL, *p <* 0.0001; two-way ANOVA) **(Figure 4D)** with significantly reduced DPOAE amplitude only at 32 kHz (32 kHz, *p* < 0.0001, two-tailed, unpaired *t*-test) **(Figure 4E & F)**. While young adult *Ocm* KO OHCs were retained **(Figure 4G & H)**, aged *Ocm* KO OHCs had a frequency dependent increase in OHC loss with the most the most significant degeneration observed at 22 kHz (*Ocm* WT = 0.333 ± 0.577%, *Ocm* KO = 40.333 ± 13.013%), 32 kHz (*Ocm* WT = 0.000 ± 0.000%, *Ocm* KO = 82.000 ± 9.626%), and 45 kHz (*Ocm* WT = 0.000 ± 0.000%, *Ocm* KO = 96.000 ± 1.826%; *p <* 0.0001, two-way ANOVA) (Figure 4I & J).

**Figure 4.**
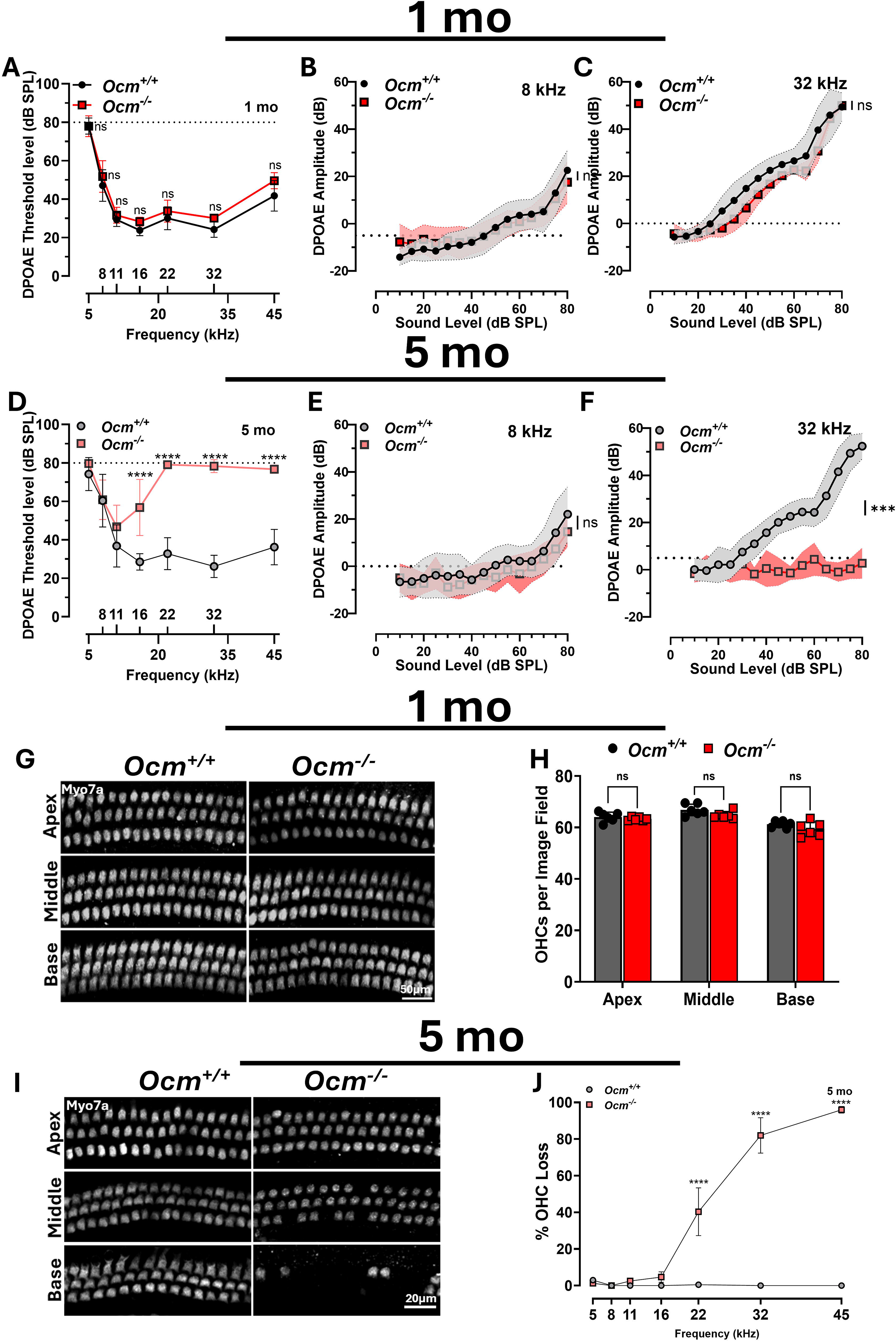
DPOAE responses and OHC integrity change with age in *Ocm* KO mice. (A –. **C**) DPOAE threshold levels **(A)** and I/O functions at f2 frequencies of 8 kHz **(B)** and 32 kHz **(C)** in young adult *Ocm* WT and KO mice. **(D – F)** DPOAE threshold levels **(D)** and I/O functions at f2 frequencies of 8 kHz **(E)** and 32 kHz **(F)** in aged *Ocm* WT and KO mice. A *2f1-f2* emission was considered present when its amplitude was above the dotted line in each graph. Data are presented as mean ± SD, n = 7 to 11 mice per experimental group. **(G & H)** Representative images of OHCs labeled with Myo7a (white) **(G)** and quantification **(H)** in the apical (5 – 11 kHz), middle (16 – 22 kHz) and basal (32 – 45 kHz) frequency regions in young adult *Ocm* WT and KO cochleae. **(I & J)** Representative images showing OHCs labeled with Myo7a (white) **(I)** and quantification **(J)** in the apical (5 – 11 kHz), middle (16 – 22 kHz) and basal (32 – 45 kHz) frequency regions in aged *Ocm* WT and KO cochleae. Statistical analysis was determined using two-way ANOVA with Bonferroni’s post hoc **(A, D, H & J)** and unpaired *t*-test **(B, C, E & F)**. *** *p < 0.001;* **** *p < 0.0001*; ns, not significant

Again, these findings suggest that the loss of OCM in OHCs precedes any changes in auditory function or sensory cell loss and that the cochlear innate immunity is sensitive to cellular dysfunction before the onset of cellular degeneration. These findings support the hypothesis that impaired calcium homeostasis in OHCs is sufficient to alter cochlear innate immunity independent of sensory cell degeneration.

### Absence of OCM has differing effects on cytokines and chemokines in the cochleae

In addition to their phagocytic function, macrophages, as mediators of innate immunity, express inflammatory receptors which influence the production of pro-inflammatory or anti-inflammatory cytokines (Lendeckel *et al*., 2022). Increased expression of inflammatory receptors and production of pro-inflammatory cytokines have been described after cochlear injury and coincide with macrophage proliferation (Fujioka *et al*., 2006; Wakabayashi *et al*., 2010; Cai *et al*., 2014; Vethanayagam *et al*., 2016). The increase in cochlear resident macrophages and alterations in morphology in the absence of physiological changes or OHC and synaptic loss, in young adult *Ocm* KO mice, prompted us to consider whether the absence of OCM also alters the expression of key innate immune signaling genes.

We performed a cytokine array on whole cochlear lysates from young adult *Ocm* WT and KO mice **(Figure 5A)**. Of the cytokines expressed on the array, we focused our quantitative analysis on pro-inflammatory and anti-inflammatory cytokines, as well as chemokines involved in macrophage recruitment **(Figure 5B – D)**. While there were no significant differences (p > 0.05, two-way ANOVA), we noticed a trend towards reduced protein expression of the pro-inflammatory cytokine, IL-6 **(Figure 5B)**. Furthermore, there was a trend towards reduced expression of the chemokine, CXCL13 **(Figure 5C)**, and the anti-inflammatory cytokines, IL-10 and IL-4 **(Figure 5D**). We also examined the mRNA expression levels of well-studied pro-inflammatory cytokines, *Il-6* and *Il-1*β, and anti-inflammatory cytokines, *Il-4* and *Il-10*. The expression level of *Il-6* was significantly lower in young adult *Ocm* KO cochlea (*Ocm* WT = 1.047 ± 0.353, *Ocm* KO = 0.353 ± 0.201, *p <* 0.05 two-tailed, unpaired *t*-test) **(Figure 5E)**. The opposite trend was observed for *Il-1*β expression with its expression levels being significantly higher in *Ocm* KO cochlea (*Ocm* WT = 1.040 ± 0.360, *Ocm* KO = 3.363 ± 0.730, *p <* 0.05 two-tailed, unpaired *t*-test) **(Figure 5F)**. While there was no significant difference in the expression level of the anti-inflammatory cytokine, *Il-4,* between *Ocm* WT and KO cochlea (*Ocm* WT = 1.350 ± 0.626, *Ocm* KO = 1.557 ± 0.669, *p >* 0.05 two-tailed, unpaired *t*-test), the expression level of *Il-10* was significantly lower in *Ocm* KO cochlea (*Ocm* WT = 1.033 ± 0.319, *Ocm* KO = 0.1300 ± 0.095, *p <* 0.05 two-tailed, unpaired *t*-test) **(Figure 5G & H)**.

**Figure 5.**
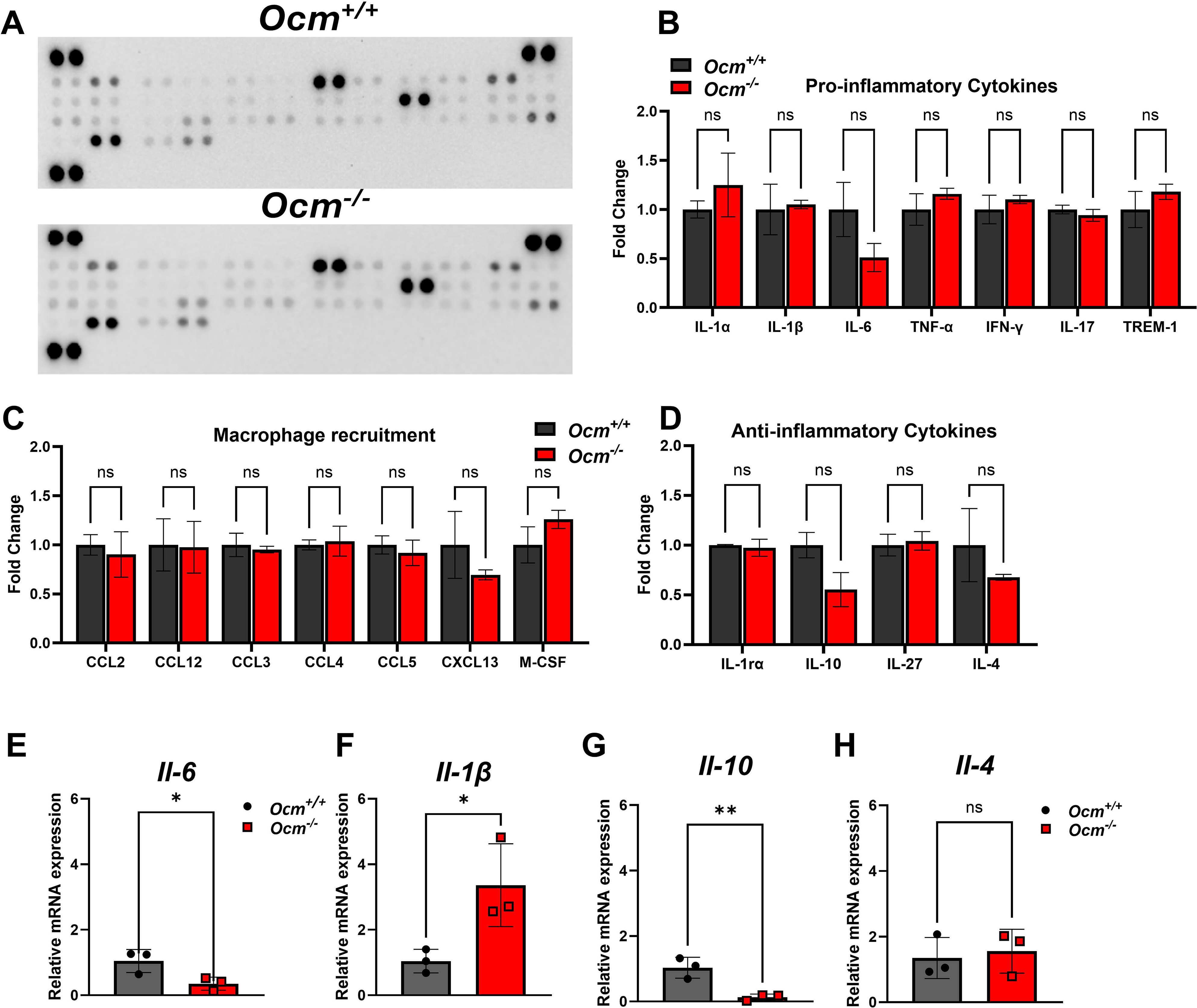
The absence of OCM leads to modest changes in inflammation-related proteins and genes. **(A)** Cytokine array blots detecting multiple cytokines and chemokines in whole cochlear lysates of *Ocm* WT and KO mice. **(B – D)** Quantitative analysis of pro-inflammatory cytokines **(B),** chemokines associated with macrophage recruitment **(C)**, and anti-inflammatory cytokines **(D)**. Relative mRNA expression levels of *Il-6* **(E)**, *Il-1*β **(F)**, *Il-4* **(G)**, and *Il-10* **(H)** in *Ocm* WT and KO cochleae. Data are presented as mean ± SD, n ≥ 2 per experimental group. Statistical analysis was determined using unpaired *t-*test * *p < 0.05;* ** *p < 0.01;* ns, not significant.

Our analysis suggests that while the absence of OCM affects the density and morphology of macrophages in the young adult cochlea, it has a variable impact on pro- and anti-inflammatory genes and proteins in the cochlea.

### Immune cell numbers are unaltered during development in *Ocm* KO mice

Cochlear macrophages are key contributors to developmental processes necessary for normal hearing onset including greater epithelial ridge (GER) remodeling, glial cell elimination and synaptic pruning, where they perform phagocytic functions (Brown *et al*., 2017; Borse *et al*., 2021; Yu *et al*., 2021; Song *et al*., 2022). Consistent with these roles, macrophage numbers gradually increase during the first postnatal week and decrease after hearing onset (P12-P14) (Brown *et al*., 2017; Dong *et al*., 2018; Miwa *et al*., 2024). One explanation for the altered innate immunity in the young adult *Ocm* KO cochlea is that it arises during cochlear development and persists into adulthood. We quantified immune cell numbers in the OSL and GER, at two developmental time periods: before hearing onset (P4 – P5) and at hearing onset (P12).

We used the progression of GER remodeling; a process associated with regulated cell death and immune cell recruitment as a proxy for assessing developmental immune changes in *Ocm* WT and KO cochleae. GER regression appeared to progress normally in *Ocm* WT and KO cochleae with the appearance of cleaved caspase 3 (apoptotic marker) positive cells in the basal region at P5 **(Figure 6A)** with near complete regression and lack of apoptotic cells at P12 **(Figure 6B)**. At P5, immune cell numbers in the apical and middle frequency regions of both *Ocm* WT and KO were comparable (apex: *Ocm* WT = 24.082 ± 6.340, *Ocm* KO = 25.105 ± 3.878, middle: *Ocm* WT = 26.833 ± 7.210, *Ocm* KO = 27.792 ± 8.518, *p* > 0.999, one-way ANOVA). Similar to previous studies, immune cell numbers significantly increased at P12 in *Ocm* WT and KO cochlea in the apex (*Ocm* WT : P12 = 36.280 ± 3.714 vs P5 = 24.082 ± 6.340 ; *Ocm* KO : P12 = 36.642 ± 6.900 vs P5 = 25.105 ± 3.878, *p <* 0.05, One-way ANOVA) and middle (*Ocm* WT : P12 = 41.255 ± 3.023 vs P5 = 26.833 ± 7.210 ; *Ocm* KO : P12 = 43.710 ± 7.904 vs P5 = 27.792 ± 8.518, *p <* 0.05, one-way ANOVA) frequency regions with no significant differences in immune cell numbers between WT and KO cochleae at P12 (*p* > 0.999, one-way ANOVA **(Figure 6C & D)**. In the basal region of *Ocm* KO cochleae at P5, there was a trend towards higher immune cell numbers compared to WT cochleae, but this was not statistically significant (*p* > 0.999, one-way ANOVA). While there was a significant increase in immune cell numbers at P12 in *Ocm* WT cochleae (P12 = 41.525 ± 3.001 vs P5 = 25.922 ± 4.817, *p* < 0.001, one-way ANOVA), *Ocm* KO cochlea did not have a significant increase (*p* > 0.999, one-way ANOVA). However, there was no significant difference between P12 immune cell numbers of *Ocm* WT and KO cochleae (*p* > 0.999, one-way ANOVA) **(Figure 6E)**.

**Figure 6.**
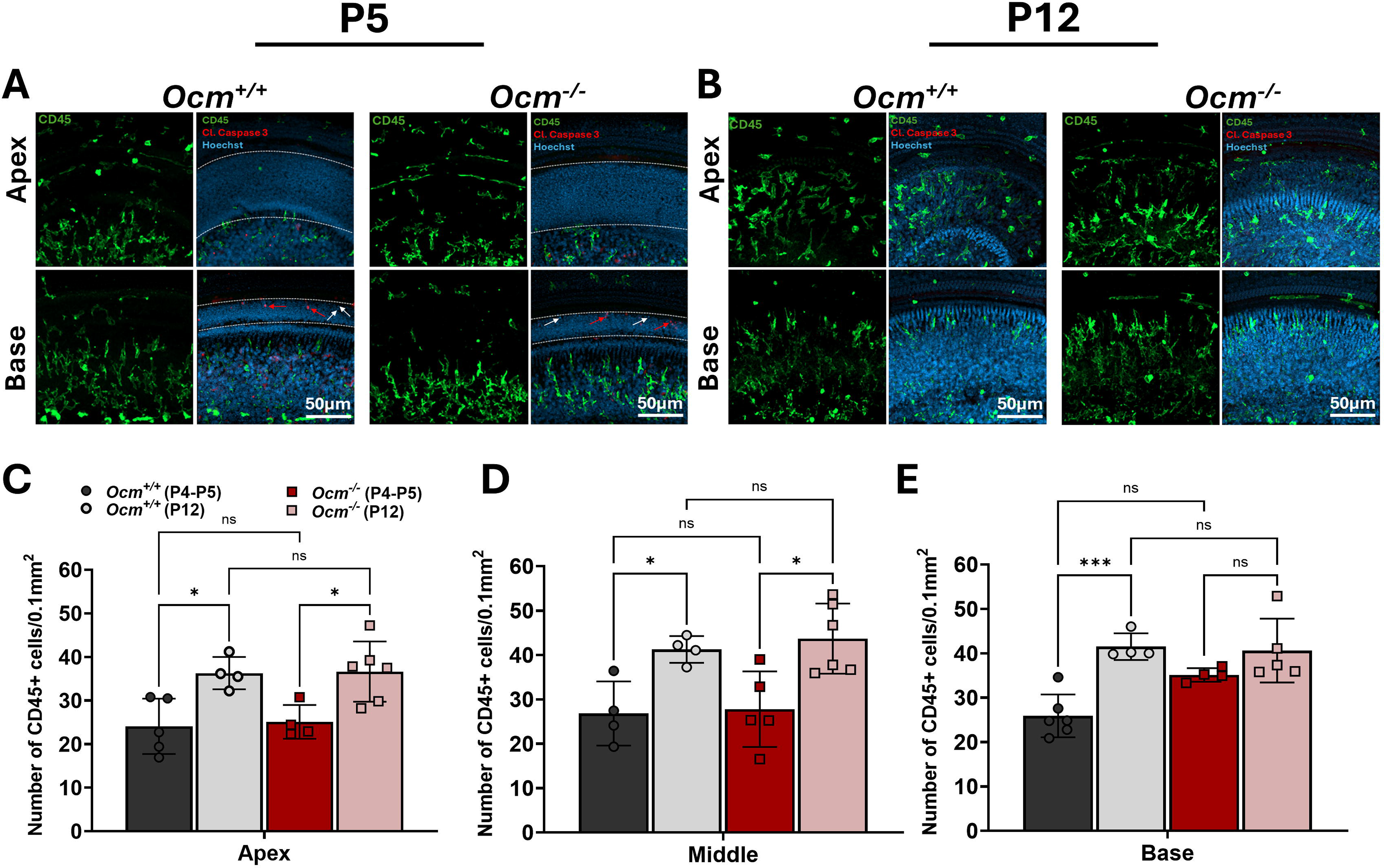
The absence of OCM has no effect on immune cell density during development. (A, **B)** Representative images of CD45 positive immune cells (green) and cleaved caspase 3 positive cells (red) in *Ocm* WT and KO cochleae at P5 **(A)** and P12 **(B)**. The white arrows point to cleaved caspase 3 positive cells. **(C – E)** Quantification of immune cell numbers of P5 and P12 *Ocm* WT and KO cochleae in the apex **(C)**, middle **(D)** and basal **(E)** frequency regions. Data are presented as mean ± SD, n ≥ 4 cochleae per experimental group. Statistical analysis was determined using one-way ANOVA with uncorrected Dunn’s test * *p < 0.05;* *** *p < 0.001;* ns, not significant.

These findings indicate that changes in macrophage density follow the normal developmental pattern in *Ocm* KO cochleae. Thus, the altered cochlear innate immunity seen in young adult *Ocm* KO cochleae is not a result of developmental differences but occurs after hearing onset when OHCs functionally mature.

## Discussion

The innate immune response of the cochlea to injury or aging is characterized by the activation and proliferation of cochlear resident macrophages as well as an increased expression of pro-inflammatory genes and proteins (Hirose *et al*., 2005; Kaur *et al*., 2015a; Frye *et al*., 2017; Frye *et al*., 2018; Zhang *et al*., 2021). In addition, certain cochlear structures such as sensory hair cells and neighboring supporting cells express proteins that mediate the cochlea’s innate immune response (Fujioka *et al*., 2006; Cai *et al*., 2014). However, the mechanisms that govern cochlear innate immunity in the absence of damage are poorly understood. In this study, we used a mouse model that lacks an OHC-specific calcium buffer, OCM, to address whether alterations in sensory cell function are sufficient to modify cochlear innate immunity.

Our findings reveal changes in macrophage density and morphology in young adult *Ocm* KO mice independent of changes to auditory function. Young adult *Ocm* WT and KO mice had comparable ABR and DPOAE threshold levels with similar DPOAE amplitudes as well as ABR wave I amplitudes and latencies. After pathological insults such as ototoxicity or noise, or aging, which are all associated with structural degeneration, macrophages proliferate (Schuknecht *et al*., 1974; Wu *et al*., 2020; Zhang *et al*., 2020). While macrophages present in the OSL of *Ocm* KO cochlea exhibited changes in density and morphology, there was no evidence of OHC or synaptic loss. After the onset of hearing loss in older *Ocm* KO mice, which is accompanied by synaptic and OHC loss, macrophage numbers remained unchanged.

In addition to increased numbers, OSL resident macrophages in *Ocm* KO mice showed greater spanned area and ramification index across all frequency regions. In microglia, these changes are associated with a surveillance phenotype, whereas an amoeboid phenotype exhibiting less ramification and spanned area is linked to activation and tissue damage (Stence *et al*., 2001; Avignone *et al*., 2015; Vidal-Itriago *et al*., 2022). In the young adult *Ocm* KO cochlea, we observed a distinct spatial partitioning of macrophages, with a uniform arrangement closer to the sensory epithelium and a more random distribution closer to the spiral ganglion. Previous reports suggest that macrophages in the OSL are arranged in roughly two rows with one row near the spiral ganglion neurons and the other row near the edge of the OSL closer to the sensory epithelium (Hu *et al*., 2018; Zhang *et al*., 2021). This arrangement might imply distinct functions for these macrophages with those near the sensory epithelium playing a role in synaptic repair or synaptic loss, and macrophages near the spiral ganglion playing a role in their survival or degeneration, after cochlear injury. However, this idea has not been systematically investigated. In our study, macrophages closer to the sensory epithelium were more ramified, consistent with a surveillance phenotype, suggesting heightened monitoring or interaction with neural elements which are particularly sensitive to injury.

While there is not a direct correlation between immune cell morphology and function, some studies have linked morphological changes to functional readouts such as increased phagocytosis and cytokine expression in the central nervous system (CNS) (Gan *et al*., 2025). A recent study in the cochlea showed that macrophages with impaired CX3CR1 signaling display an amoeboid morphology, which correlated with elevated CD68 expression, a phagocytic marker, under both naïve and noise-damaged conditions (Gawande *et al*., 2025). Another study showed that CD68 expression varied across cochlear regions after treatment with the ototoxic compound, cyclodextrin, but failed to assess macrophage morphology in these regions (Ye *et al*., 2025). In the present study, while we observed morphological changes associated with a surveillance phenotype in *Ocm* KO cochleae, OSL macrophages expressed CD68 in both *Ocm* WT and KO cochleae. However, there was greater expression of CD68 in *Ocm* KO macrophages in basal regions. The variable expression of CD68 coupled with the increased surveillance morphology seen in *Ocm* KO OSL macrophages suggest that there is not a clear link between macrophage morphology and the expression of activation markers such as CD68. To better understand changes in the function of OSL macrophages in *Ocm* KO cochleae, future studies could focus on incorporating macrophage phagocytic assays in conjunction with multiple cell surface markers to determine the activation states of *Ocm* KO macrophages. Collectively, while there were changes in the morphology of OSL macrophages representing a shift in baseline immune homeostasis, we cannot conclusively infer the function of these macrophages.

To determine whether the changes in macrophage numbers, organization and morphology in *Ocm* KO mice accompanied changes in genes associated with innate immune signaling, we assessed protein and mRNA expression levels of cytokines and chemokines. While we observed trends toward decreased protein levels of the pro-inflammatory cytokine, IL6, and anti-inflammatory cytokines, IL-4 and IL-10 in *Ocm* KO cochlea, these results did not reach significance. Gene expression analysis of the pro-inflammatory cytokines *Il-6* and *Il-1*β showed unusual expression patterns. In the cochlea, the levels of both these cytokines increase after ototoxic and acoustic injury along with an increase in immune cells (Satoh *et al*., 2003; Hashimoto *et al*., 2004; Fujioka *et al*., 2006; So *et al*., 2007; Tan *et al*., 2016). In the present study, there was an increased expression of *Il-1*β without a corresponding increase in *Il-6*. Several studies in the cochlea and other disease models show that the levels of these pro-inflammatory cytokines often increase concurrently. However, studies by Tan *et al.,*(2016) and Fujioka *et al.,*(2006) suggest an earlier increase of *Il-1*β before a corresponding increase in *Il-6* after noise exposure mirroring in vitro studies (Ichimiya *et al*., 2003). Alternatively, the increased expression of *Il-1*β without a corresponding increase in *Il-6*, in *Ocm* KO cochleae, might suggest a state of low-grade inflammation. Similar to the pro-inflammatory cytokines, the anti-inflammatory cytokines, *Il-4* and *Il-10*, exhibited unusual expression levels. These cytokines typically play roles in macrophage polarization to a reparative phenotype. While there was no significant difference in *Il-4* expression between *Ocm* WT and KO cochleae, *Il-10* expression was significantly lower in *Ocm* KO cochleae compared to WT. Previous studies in the cochleae, *Il-10* deficiency is associated with exacerbated hearing loss and increased recruitment and proliferation of reactive immune cells in models of autoimmune hearing loss and inflammation-mediated cochlear damage (Woo *et al*., 2015; Mwangi *et al*., 2017). Taken together, the increased density of macrophages and surveillance morphology seen in post-hearing *Ocm* KO mice as well as the varying changes in cytokines cannot be attributed to hearing loss or degeneration of cochlear structures. Although, these changes might be associated with the earlier onset of progressive hearing loss and susceptibility to noise trauma in *Ocm* KO cochleae.

Purinergic signaling plays a major role in shaping inflammatory responses. ATP released from stressed or damaged cells can initiate pro-inflammatory cytokine release via P2X7 receptors and can bind P2X receptors on macrophages further exacerbating inflammatory responses (Burnstock, 2016; Liu *et al*., 2023). Because calcium dysregulation can influence ATP and downstream purinergic signaling, both of which are processes that mediate macrophage function (Desai & Leitinger, 2014; Entsie et al., 2023), changes in OHC calcium buffering may have an indirect effect on shaping cochlear immune homeostasis. Our lab has previously reported increased gene and protein expression of purinergic receptors in the cochlea of *Ocm* KO mice compared to WT mice (Yang *et al*., 2023; Yang *et al*., 2026). Additional studies in the young adult show that after prolonged noise exposure (95 dB SPL, 9 hrs), *Ocm* KO mice retained their already elevated baseline P2X2 levels while WT mice showed the expected noise-induced upregulation of P2X2 (Yang *et al*., 2026), consistent with previous studies (Wang *et al*., 2003). In the CNS, several studies have demonstrated increased expression of P2X receptors in microglia following injury and these changes correlate with morphological alterations and functional activation, including phagocytosis (Ulmann *et al*., 2008; Burnstock, 2016). (Ni *et al*., 2013) showed that silencing P2X7 enhanced phagocytosis of amyloid-β plaques by microglia, suggesting the context-dependent nature of purinergic signaling. Although purinergic receptor expression levels were not assessed in this study the role of P2X receptors in shaping immune responses combined with the elevated P2X receptor levels seen in *Ocm* KO mice raises the possibility that altered ATP-P2X signaling may contribute to the immune phenotypes we see in *Ocm* KO mice.

To determine if the absence of OCM also alters cochlear innate immune homeostasis during development, we quantified immune cell numbers at two developmental timepoints – P4 – P5, prior to hearing onset, and P12, the onset of hearing. Macrophages populate the otocyst beginning around embryonic day 10 (Hirose *et al*., 2017; Kishimoto *et al*., 2019) with their numbers increasing during postnatal maturation. The timing of this increase corresponds to developmental processes such as glial cell elimination, synaptic pruning and GER regression, all of which are marked by increased cell death (Dong *et al*., 2018; Yu *et al*., 2021; Song *et al*., 2022; Miwa *et al*., 2024). During these processes, macrophages are believed to play phagocytic roles, and the depletion of cochlear macrophages results in impaired auditory function (Brown *et al*., 2017). As the cochlea matures, macrophage numbers decrease across all cochlear regions and baseline levels are achieved around P17 – P21 (Dong *et al*., 2018). In the present study, immune cell numbers followed expected developmental trends, with the highest numbers observed at P12 in both *Ocm* WT and KO. We found no significant differences in immune cell numbers between *Ocm* WT and KO mice at either P5 or P12 across all frequency regions. These findings suggest that the absence of OCM does not alter cochlear innate immune homeostasis prior to, or at the onset of hearing, but after hearing onset, coinciding with the functional maturation of OHCs.

In summary, our findings demonstrate that changes in OHC calcium homeostasis can influence cochlear innate immune cells prior to auditory decline. The lack of OCM altered the cochlear innate immune homeostasis marked by increased OSL macrophage numbers and morphological changes associated with heightened surveillance. Together with previous evidence of increased purinergic receptor expression and dysfunctional mitochondria in *Ocm* KO OHCs, our data supports a model whereby impaired OHC function at the cellular level induces sustained changes in OSL macrophages that may influence long-term cochlear vulnerability. These results reveal a link between OHC calcium homeostasis and cochlear innate immune regulation, and that the cochlear immune system may be capable of sensing impaired OHC cellular function before the onset of hearing loss or degeneration of cochlear structures.

## CRediT authorship contribution statement

**Weintari D. Sese**: Conceptualization, Formal analysis, Investigation, Methodology, Software, Visualization, Data curation, Writing – Original Draft, Writing – Review & Editing. **Janith N. Halpage**: Investigation, Writing – Review & Editing. **Mahika V. Palani**: Investigation, Formal Analysis, Writing – Review & Editing. **Evan J. Paltjon:** Investigation, Writing – Review & Editing. **Kiah C. Sleiman**: Investigation, Writing – Review & Editing. **Aubrey J. Hornak**: Writing – Review & Editing, Resources, Supervision, Project administration. **Dwayne D. Simmons**: Conceptualization, Methodology, Formal analysis, Investigation, Resources, Data Curation, Writing – Original Draft, Writing – Review & Editing, Visualization, Validation, Supervision, Project administration, Funding acquisition.

## Funding

Research was supported by the National Institute on Deafness and Other Communication Disorders of the National Institutes of Health under award number 1R15DC022073-01 (Dwayne D. Simmons).

**Figure S1. Macrophages in the OSL of oncomodulin KO mice exhibit distinct spatial clustering. (A, D)** Binarized images of the spatial distribution of OSL macrophages in the apex **(A)** and base **(D)** of *Ocm* KO cochleae. **(B, E)** Scatter plot of individual macrophage centroids mapped by X (horizontal) and Y (vertical) coordinates. Each dot represents the centroid of a macrophage in the image field **(C, F)** Y centroid values of macrophages assigned to cluster 1 and cluster 2. Lower Y values indicate cells nearer to the top of the image field. Results reported are from a single representative image. Data are presented as mean ± SD, two-way ANOVA with Bonferroni’s post hoc. **** *p < 0.0001*.

