## Supplementary figures and images for "Cochlear Innate Immune Homeostasis is altered in the Oncomodulin-Deficient Mouse Model"

### Supplementary Figure 1

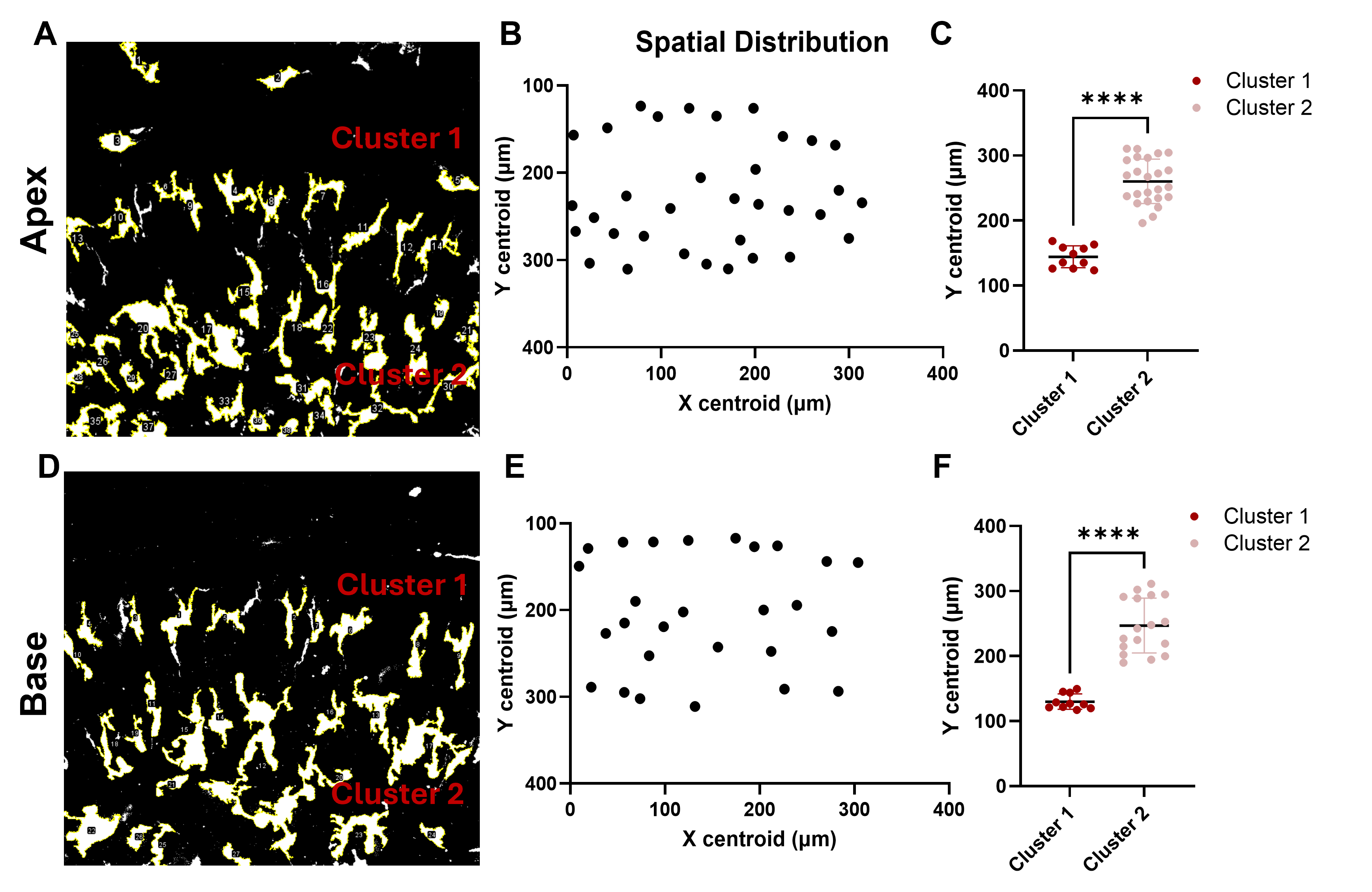
